# Glucocorticoid Regulation of Ependymal Glia and Regenerative Potential after Spinal Cord Injury

**DOI:** 10.1101/328781

**Authors:** Craig M. Nelson, Han Lee, Randall G. Krug, Aichurok Kamalova, Nicolas N. Madigan, Karl J. Clark, Vanda A. Lennon, Anthony J. Windebank, John R. Henley

**Affiliations:** Departments of Neurological Surgery, College of Medicine, Rochester, MN, USA 55905.; Departments of Biochemistry and Molecular Biology, College of Medicine, Rochester, MN, USA 55905.; Departments of Neurology, College of Medicine, Rochester, MN, USA 55905.; Departments of Laboratory Medicine and Pathology, College of Medicine, Rochester, MN, USA 55905.; Departments of Immunology, College of Medicine, Rochester, MN, USA 55905.; Departments of Physiology and Biomedical Engineering, College of Medicine, Rochester, MN, USA 55905.; Departments of Mayo Graduate School, Mayo Clinic, College of Medicine, Rochester, MN, USA 55905.

**Keywords:** Neural regeneration, spinal cord injury, glucocorticoid signaling, Nr3c1, ependymalglia, glial bridge

## Abstract

Following injury, the mammalian spinal cord forms a glial scar and fails to regenerate. In contrast, spinal cord tissue of vertebrate fish regenerates and restores function. Cord transection in zebrafish (*Danio rerio*) initially causes paralysis and neural cell death, with subsequent ependymal glial proliferation, extension of bipolar glia across the lesion, and neurogenesis. Axons extending from spared and nascent neurons along trans-lesional glial bridges restore functional connectivity. Here we report that glucocorticoids directly target the regeneration supporting changes in ependymal glia to inhibit neural repair. This effect is independent of hematogenic immune cells or microglia. Furthermore, glucocorticoid receptor signaling in ependymal glia is inversely regulated in rat models of spinal cord injury compared to zebrafish. The blockade of neural regeneration by glucocorticoids via a direct effect on ependymal glia has important clinical implications concerning the putative therapeutic benefit of corticosteroids in early management of spinal cord injury.

## Introduction

Spinal cord injury causes life-long physical disability. The goal of contemporary treatment is to limit morbidity by stabilizing the injury and managing inflammation, but restoration of function is unattainable (1–3). For many decades glucocorticoid (GC) therapy has been considered a means for limiting tissue damage and functional loss after spinal cord injury (1, 2, 4). Re-evaluation of evidence from this early work revealed study design limitations, and subsequent *post hoc* analyses failed to validate improvements in the critical primary outcome measures of motor and sensory function (5–8). The inconsistency in reported results and significant systemic complications have led surgical societies to downgrade the recommendation of GC therapy from the previous *standard-of-care* for medical practice (highest scientific validity) to an *option* (lowest validity) (9, 10). Nevertheless, spine surgeons continue to employ GCs in treating spinal cord injuries (3). Mechanistic investigation is needed to resolve this controversial issue.

Lack of significant regeneration after mammalian cord injury is commonly attributed to spinal neuron loss, quiescence of spared cells, inflammation, and a non-permissive microenvironment for axon regrowth. Despite evidence that astroglia can support regeneration (11), the scar formed by reactive astroglial proliferation is considered a barrier to axon regrowth and functional recovery (12–14). The nonpermissive environment of the injury site limits interpretation of potential positive and negative regulators of mammalian cord regeneration.

As a complementary approach, comparison of signaling events in a regeneration-permissive vertebrate model holds potential to unmask differences in the mammalian signaling pathways induced by spinal cord injury. The zebrafish is a powerful model for identifying novel molecular and cellular mechanisms that regulate neural repair because its spinal cord regenerates functionally following complete transection (15, 16) This is in part due to re-entry of ependymal glia, a type of radial glia, into the cell cycle. These cells also form bipolar bridges that span the lesion to support axon regeneration (17, 18), and yield multipotent neural precursor cells that replenish lost neural cell types (19, 20).

In zebrafish models of acute CNS injury (both adult brain injury [21] and transecting spinal cord injury [22]), administration of GCs suppressed immune cell infiltration and this correlated with reduction of radial glial proliferation and regeneration. The role of GCs in regulating CNS neurogenic niches is poorly defined, but they have been reported to attenuate hippocampal neurogenesis in mammals and regeneration competence in the chick retina (23–26).

A prominent GC receptor that is common to mammals and zebrafish is Nuclear receptor subfamily 3, group C, member 1 (Nr3c1, 27, 28). When ligated by GC, this intracellular receptor enters the nucleus and serves as a transcription factor (29). Its binding to DNA encoding GC response element sequences stimulates transcription of target genes (30), particularly those promoting anti-inflammatory pathways (31). Nr3c1 is expressed widely in the adult rat spinal cord and increases within 4 h of spinal cord injury (32). Yet the mechanistic actions of GCs on neural cells during functional regeneration after cord injury remain undetermined.

While investigating mechanisms underlying functional regeneration of the injured spinal cord in larval zebrafish, we discovered that GCs inhibit neural repair by targeting Nr3c1-expressing ependymal glia. Comparative investigation in adult rats revealed that the signaling activity of this receptor in ependymal glia was inversely regulated following complete spinal cord transection. These complementary data plausibly explain the limited neural repair that occurs in the mammalian CNS, and suggest that conventionally-administered high-dose corticosteroid therapy likely compounds failure of the injured spinal cord to regenerate.

## Results

### Ependymal glial Nr3c1 is inversely regulated in zebrafish and rat after spinal cord injury

Regeneration of the injured spinal cord in zebrafish is potentially regulated by loss of the inhibitory signals that normally constrain regeneration-competent cells in a latent state. Based on earlier studies in zebrafish (21, 22) and chicks (25, 26), we hypothesized that GCs were an inhibitory factor. We therefore sought to identify and compare the cell types targeted by GCs after spinal cord injury in the robust regenerative model of larval zebrafish and the non-permissive milieu of adult rats.

Although recognized to be expressed widely (32), little is known of the activity of the neural GC receptor Nr3c1 in spinal cord regeneration. To identify the cellular targets of GCs and investigate the potential role of Nr3c1 in neural repair, we immunostained neural tissue adjacent to a cord lesion (Fig. 1A-C). In non-injured control fish, Nr3c1 immunoreactivity was strong in nuclei of ependymal glia (positive for glial fibrillary acidic protein [Gfap,]) surrounding the central canal (Fig. 1A). The nuclear Nr3c1 localization was consistent with constitutive receptor stimulation. Gfap-negative cells that highly expressed nuclear Nr3c1 were identified as neurons (dual HuC/D nuclear immunoreactivity, not shown). Following spinal cord injury, Nr3c1 immunoreactivity was reduced in intensity. By 24 h post spinal cord injury, Nr3c1 was cytoplasmic in ependymal glia, and immunoreactivity was decreased further at 48 h (Fig.1 B, C). The observation of decreased expression and cytoplasmic redistribution of Nr3c1 was consistent with loss of Nr3c1 signaling following transecting injury. The dynamic Nr3c1 expression suggested that ependymal glia were a potential GC target.

**Figure 1:**
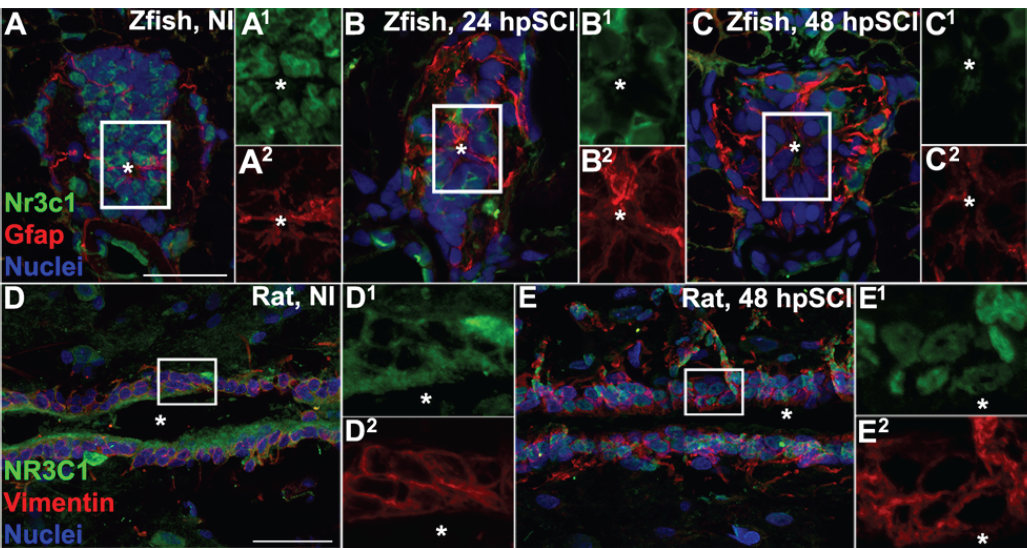
SCI Stimulates Differential Nr3c1 Expression by Ependymal Glia in Zebrafish and Rats. (A-C) Immunolabeling for Nr3c1 (green), Gfap (red) and DAPI staining (blue) in larval zebrafish spinal cord transverse sections at 24 h post no injury (A), 24 (B) and 48 (C) h post SCI. The nuclear Nr3c1 distribution in Gfap+-ependymal glia without injury (A^1, 2^) is shifted to cytoplasmic at 24 and 48 h post SCI (B^1, 2^, and C^1, 2^). Scale bar, 20 µm. Boxed regions denote ependymal glial cell bodies around the central canal and are magnified 167% (A^1, 2^, B^1, 2^, and C^1, 2^). Representative images from n = 10 zebrafish (3 sections per cord examined). See supporting Fig. S1. (D and E) Adult rat spinal cord longitudinal sections examined at 48 h post no injury (D) and 48 h post SCI (E). The cytoplasmic Nr3c1 expression in ependymal glia in controls (D^1, 2^) is shifted to the nucleus after SCI (E^1, 2^). Scale bar, 50 µm. Boxed regions denote ependymal glial cell bodies around the central canal and are magnified 310% (D^1, 2^ and E^1, 2^). Representative images from n = 3 rat spinal cords (3 sections per cord examined).

Comparative immunohistochemical staining of non-injured rat spinal cord revealed NR3C1 immunoreactivity co-expressed with vimentin in ependymal glia surrounding the central canal (Fig. 1 D, E). By 48 h post spinal cord injury, NR3C1 expression in rat ependymal glia had dramatically redistributed from the cytoplasm to the nuclear compartment. The nuclear translocation of NR3C1 indicated receptor activation. Thus, NR3C1 distribution and expression are differentially regulated in ependymal glia after spinal cord injury in rats compared to zebrafish. NR3C1 expression in ependymal glia implicates a direct effect of GCs on these neural cells.

### Glucocorticoids block functional recovery from spinal cord injury via Nr3c1

Finding that Nr3c1 expression is differentially regulated following spinal cord injury in larval zebrafish and adult rats led us to investigate whether GC pathway activation might be sufficient to block functional regeneration. In larval zebrafish, functional recovery from spinal cord injury is robust by 120 h after transection (33, 34). We used a quantitative locomotor recovery assay (17) to evaluate functional improvement of swimming ability as an outcome measure after complete spinal cord transection. Compared to non-injured control fish, the swimming response was severely impaired at 6 h, but had improved by 24 and 48 h after transection (Fig. 2). Swimming behavior was robust by 72 h and nearly indistinguishable from that of non-injured groups by 120 h. In contrast, locomotor recovery was significantly less in fish treated with the synthetic glucocorticoid Dexamethasone (Dex; *p* < 0.01). Dex treatment without spinal cord injury did not significantly impair locomotor responses (*p* = 0.15). We next mutated *nr3c1* by customized genome editing using transcription activator-like nucleases (TALENs) and compared the effect of non-targeting GM2 TALENs. RT-PCR confirmed *nr3c1* mutation in embryos injected with mRNAs encoding targeting TALENs (46; not shown), and immunostaining verified that spinal cord Nr3c1 expression was reduced compared to in GM2 controls (Supplemental Fig. S1). Locomotor recovery following spinal cord injury and Dex treatment in fish with mutated *nr3c1* did not significantly differ from controls at 120 h post spinal cord injury (*p* = 0.85), and the *nr3c1* TALEN injected zebrafish controls (no injury, no Dex) had no behavioral phenotype compared to non-transected controls (*p*= 0.36). Thus, inhibition of locomotor recovery from spinal cord injury by GCs is Nr3c1 dependent.

**Figure 2:**
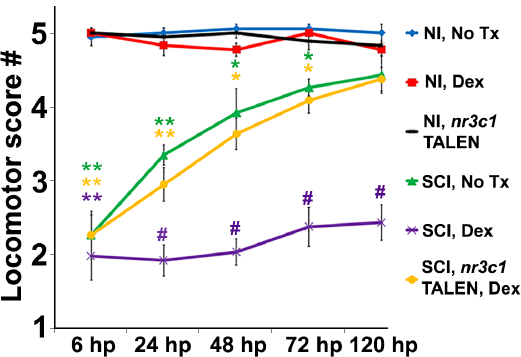
Glucocorticoids Inhibit Functional Recovery following SCI. Locomotor recovery score of wild-type or *nr3c1* targeting TALEN-injected zebrafish that received no treatment or Dex at 6, 24, 48, 72 and 120 h post no injury or SCI. Locomotor responses were quantified from no injury with no treatment controls (blue line), Dex (red line), and *nr3c1* TALEN injections (black line), or from SCI with no treatment (green line), plus Dex (purple line), and plus Dex and *nr3c1* TALEN injections combined (yellow line). Data show the mean ± SEM; *n* = 6 per condition (*, *p* < 0.05; **, *p* < 0.01 [compared to no injury], and #, *p* < 0.01 [compared to both no injury and SCI no treatment]). * and # color coded to match corresponding data points. See supporting Fig. S1.

To evaluate the extent of cell death following spinal cord injury, and to determine whether or not GCs altered cell viability (a potential reason for decreased locomotor recovery), we compared DNA fragmentation via TUNEL assays in the transected cords of zebrafish treated with Dex with cords of controls. Counts of TUNEL-positive nuclei from within 100 µm of the lesion center in whole-mount preparations showed that Dex treatment did not significantly change the number of cells dying at the peak of spinal cord injury-induced apoptosis (*p* = 0.3; Supplemental Fig. S2). We concluded from these findings that impairment of locomotor recovery in Dex recipients is attributable to regulation of a critical aspect of regeneration rather than to altered cell viability.

### Glucocorticoids inhibit regeneration-supporting ependymal glia

Earlier studies demonstrated an increased number of neurogenic ependymal glia reentering the cell cycle after cord injury (17, 19, 20, 35). However, the regulating molecular processes are not fully understood. Based on our data concerning Nr3c1 expression in the zebrafish spinal cord (Fig. 1), we hypothesized that GCs inhibit functional recovery by impairing the capacity of ependymal glia to support regeneration.

To evaluate the potential role of GCs in regulating cell division following spinal cord injury, we counted proliferating neural cells (PCNA-positive nuclei) within 100 µm of the lesion center, comparing Dex-treated and non-treated zebrafish (Fig. 3 A-D, H). Compared to no treatment, Dex did not significantly alter proliferation in non-injured controls (average PCNA-positive nuclei 12.6 ± 2.4 and 13.8. ± 2.6, respectively, *p* = 0.73). After transection, proliferation had started by 24 h (55.6 ± 11.8), was maximal at 48 h (137.6 ± 10.2), and progressively fell at 72 h (99.1 ± 8.3) and 120 h (45.0 ± 6.9). In contrast, proliferating cells in lesioned regions were significantly fewer in Dex-treated fish at all time points examined after spinal cord transection (24 h, *p* = 0.04; 48 h, *p* < 0.01; 72 h, *p* < 0.01; 120 h, *p* = 0.03). These data are consistent with GCs being inhibitory to proliferation.

**Figure 3:**
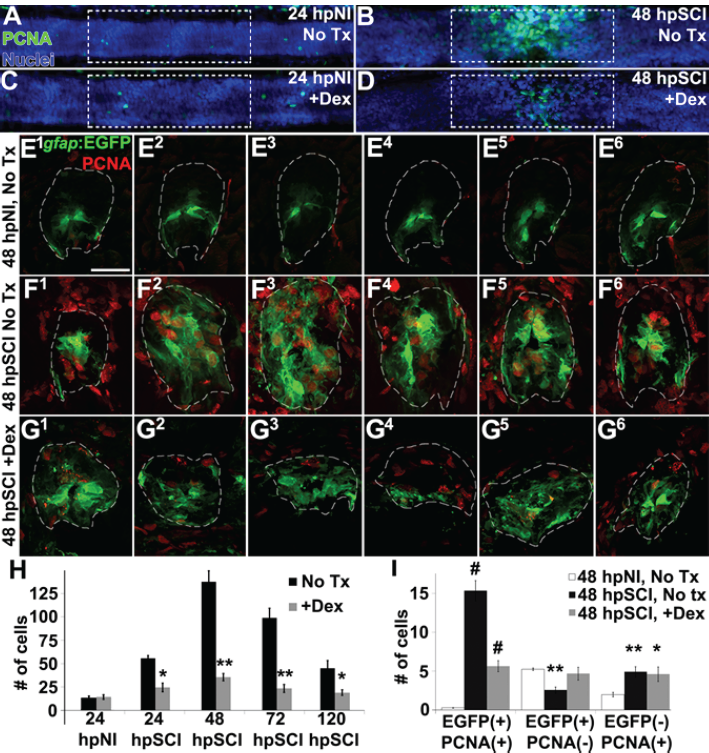
Glucocorticoids Inhibit Ependymal Glial Proliferation following SCI. (A-D) Whole-mounted larval zebrafish show immunolabeling for PCNA (green) and DAPI staining (blue) in the spinal cord after no injury (A and C) and 48 h post SCI (B and D), either followed by no treatment (A and B) or Dex treatment (C and D). Dashed boxes represent a 200-µm region centered at the lesion site used for quantifying cell counts. (E-G) Spinal cord transverse sections in Tg(*gfap*:EGFP) zebrafish were immunolabeled to detect PCNA (red) and EGFP in six concurrent 16-µm thick cryosections that spanned the lesion site at 48 h post no injury (E^1^ - E^6^), 48 h post SCI followed by no treatment (F^1^ - F^6^), and 48 h post SCI + Dex (G^1^ - G^6^). Scale bar, 20 µm (E-G). Dashed outlines define the spinal cord area used for quantifying cell counts. (H)Mean number of PCNA-positive cells ± SEM quantified from within a 100-µm region centered on the lesion (dashed boxes in A-D). *, *p* < 0.05; **, *p* < 0.01; n = 10. Comparisons were to the no treatment group for each time point. (I)Mean number of PCNA-positive and EGFP-positive cells ± SEM quantified within spinal cord (dashed outlines in E-G) cross sections in no injury controls, SCI without treatment, and SCI + Dex. *, *p* < 0.05 and **, *p* < 0.01 compared to no injury; #, *p* < 0.05 and ##, *p* < 0.01 compared to both no injury and SCI control groups; n = 6.

To identify the cell population with diminished proliferation, we evaluated transgenic reporter fish expressing EGFP in radial glia (Tg(*gfap*:EGFP;36; Fig. 3 E^1^-G^6^). We compared numbers of dual PCNA-positive and EGFP-positive glia and non-ependymal cells (EGFP-negative) in 6 consecutive sections spanning the spinal cord lesion (dashed outline) and in non-injured controls (Fig. 3I). As anticipated, the average number of proliferating ependymal glia was significantly greater in regions of transecting injury (15.6 ± 3.3) than in corresponding regions of non-injured controls (0.3 ± 0.2, *p* < 0.01), and Dex treatment significantly reduced the proliferating ependymal glial cell numbers in lesional regions (5.5 ± 1.8, *p* < 0.01). Following injury, quiescent ependymal glial cell numbers were significantly less than in non-injured controls (2.6 ± 1.0 compared to 5.3 ± 0.4, *p* < 0.01) but numbers in Dex-treated spinal cord injury groups were not significantly altered (4.6 ± 2.2, *p* = 0.55). Dividing non-ependymal cell numbers also were increased (4.3 ± 0.7) compared to non-injured controls (1.9 ± 0.7, *p* < 0.01), but that population was not significantly altered by Dex treatment (4.9 ± 2.3, *p* = 0.80). Thus, Dex treatment significantly reduced post-injury cell proliferation, of which ∼75% of PCNA-positive nuclei were EGFP-positive ependymal glia or daughter NPCs. These data agree with results reported previously from adult zebrafish studies, namely that the predominant dividing cell type in response to spinal cord injury is Tg(*gfap*:EGFP) expressing ependymal glia and their daughter NPCs (17, 19, 20, 35). We conclude from these findings that GCs inhibit ependymal glial proliferation.

Morphologically, ependymal glia in the transected zebrafish cord form elongated bipolar bridges that span the lesion site to support trans-lesion axon regeneration (17, 37). To investigate whether or not GCs regulate bipolar glial bridge formation following cord injury, we compared Dex-treated and control Tg(*gfap*:EGFP) zebrafish. Ependymal glia in non-injured cords paralleled the entire central canal (Supplemental Fig. S3). At 24 h after injury, ependymal glia adjacent to the lesion extended elongated processes into the disorganized injury site and contacted the opposing side. However, with Dex treatment, the rostral and caudal cord remained separated, and bipolar glial bridges were absent. Thus, within 24 h of transection, ependymal glia in the injured larval spinal cord become bipolar and form trans-lesion glial bridges. The blockade of glial bridge formation by Dex treatment establishes an inhibitory role for GCs.

We next investigated whether GCs might modulate axon regrowth in the context of morphological changes in ependymal glia of Tg(*gfap*:EGFP) zebrafish treated with Dex (Fig. 4). Acetylated tubulin immunoreactivity identified axons in whole-mount preparations. We measured axons and ependymal glia at the lesion site at specified time points by corrected total fluorescence (38; Fig. 4I,). Cords of non-injured control fish were densely populated with axons and ependymal glia. At 12 h after injury, the earliest time point studied, the fluorescence intensity of axons and ependymal glia was significantly reduced (quantification Fig. 4I, *p* < 0.01). At 24 h, axons within the lesion were closely associated with bipolar glial bridges, and at 48 h their fluorescence intensity had increased. By 72 h, the fluorescence intensity of axons had reached pre-injury levels, rostral and caudal sides of the cord were rejoined, and ependymal glial cells were abundant at the lesion site. By contrast, glial bridges did not form in Dex recipients and axons failed to span the lesion site at 24 h post cord transection (*p* = 0.01). At 48 h and 72 h, axon regrowth in Dex recipients remained significantly attenuated (Fig. 4I, *p* < 0.01). Thus, regeneration was initiated within 24 h of cord injury with axons associating with bipolar glial bridges spanning the lesion site, and restored to their pre-injury state by 72 h. Dex inhibited both the formation of bipolar ependymal glial bridges and subsequent axon regrowth. These findings are consistent with the notion that trans-lesion axon regrowth depends on ependymal glial bridge formation (17).

**Figure 4:**
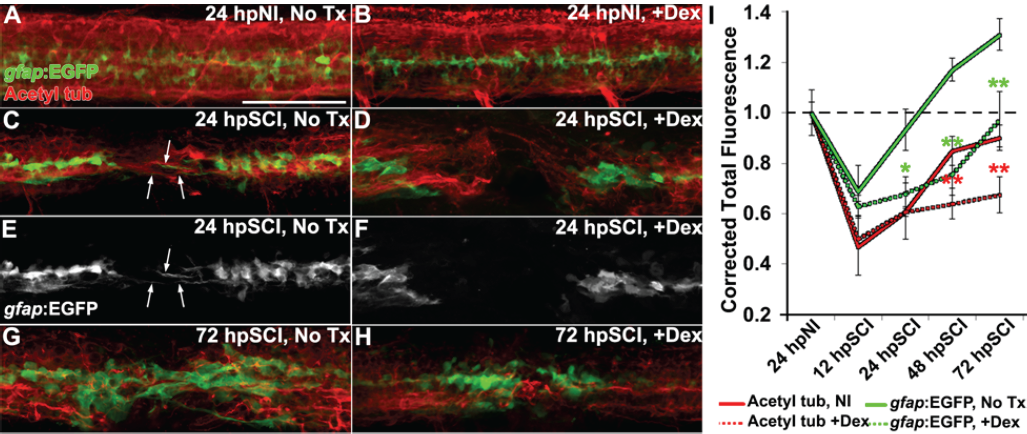
Glucocorticoids Suppress Glial Bridges and Axon Regrowth. (A-H) Whole-mounted Tg(*gfap*:EGFP) zebrafish (lateral view) show spinal cord immunolabeling for EGFP (green) and acetylated tubulin (axons, red) in non-injured controls (A and B), SCI followed by no treatment (C, E, and G), and SCI with Dex treatment (D, F, and H) for 24 and 72 h post SCI. Arrows mark bipolar EGFP-positive glial bridges (C and E) and juxtaposed axons (C) that had entered the lesion at 24 h post SCI, but were absent with Dex treatments (D and F). Scale bar, 50 µm (A-H). (I) Quantification of the mean corrected total fluorescence intensity for axons (red lines) and ependymal glia (green lines) within 50 µm of the lesion center from no treatment controls (solid lines) and +Dex (dashed lines) relative to no injury controls. Data show means ± SEM. *, *p* < 0.05; **, *p* < 0.01 compared to SCI no treatment; n = 7. * and ** color coded to match corresponding data points. See related Fig. S3.

We next assessed, by EdU uptake, the regeneration of neurons and glia in Tg(*gfap*:EGFP) zebrafish treated with Dex (Fig. 5). EdU-positive cells were quantified within 100 µm of the lesion center and identified as EGFP+ (glia), HuC/D+ (neurons), or of unknown lineage (Fig. 5 D). EdU-positive glial cell numbers were significantly increased at 120 h post transection injury by comparison to non-injured fish (*p* < 0.01). Significantly fewer EdU-positive ependymal glia were present after injury in Dex recipients (*p* <0.01). Similarly, EdU-positive neuron numbers were significantly increased after injury (*p* < 0.01), and significantly decreased after injury in Dex recipients (*p* < 0.01). Numbers of non-identifiable EdU-positive cells significantly increased following injury (*p* = 0.02), but did not change significantly with Dex treatment (*p* = 0.11). Overall, our data demonstrate that attenuation of neurogenesis by GCs following spinal cord injury is likely due to inhibition of ependymal glial proliferation (Fig. 3), which is the source of new neural cells.

**Figure 5:**
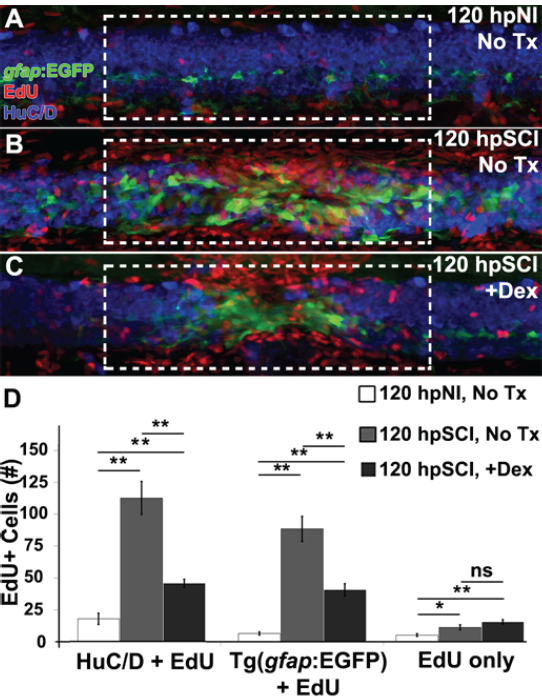
Glucocorticoids Suppress Neurogenesis. (A-C) Whole-mount preparations (lateral view) of Tg(*gfap*:EGFP) transgenic zebrafish show immunolabeling for EGFP (green), HuC/D (neurons, blue) and detection of EdU (red) from 120 h post EdU incubation and no injury (A), SCI followed by no treatment (B), and +Dex (C). Dashed boxes represent a 200-µm region centered at the lesion site used for quantifying cell counts. (D) Quantification of the mean number of EdU-positive cells within the spinal cord (± SEM). *, *p* < 0.05; **, *p* < 0.01; ns, not significant; n = 10.

### Ependymal glia are glucocorticoid targets in spinal cord injury

To investigate the potential involvement of hematopoiesis-derived cells in initiating regeneration, we utilized the bloodless *cloche* mutant zebrafish (*clo*^m39^; 39). Acting at a very early developmental stage, upstream of genes required for hematopoiesis and vasculature formation (40–44), the *cloche* mutation allows analysis of regeneration in an immune-deficient background (45). Following complete transection, whole-mount preparations of homozygous *clo*^m39^ and wild-type zebrafish spinal cords were immunolabeled for PCNA and compared (Fig. 6). At 48 h, the peak of ependymal glial proliferation (Fig. 3), injured *clo*^m39^ spinal cords contained significantly more PCNA-positive nuclei (96.8 ± 12.6) than non-injured cords (7.5 ± 2.0, *p* < 0.01), but did not differ significantly from transected wild-type cords (106.5 ± 4.6, *p* = 0.61). These results suggested that initiation of spinal cord regeneration is independent of hematopoiesis-derived cells.

**Figure 6:**
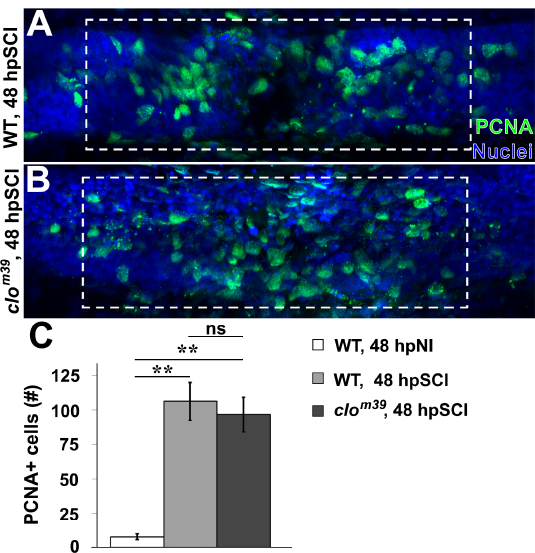
SCI-induced Proliferation Occurs without Hematogenic Cell Responses. (A and B) Whole-mount preparations (lateral view) show spinal cords of wild-type (A) and *cloche* mutant (B) zebrafish immunolabeled for PCNA (green) at 48 h post SCI. Dashed boxes represent a 200-µm region centered at the lesion site used for quantifying cell counts. (C) Quantification of PCNA-positive nuclei from within 100 µm centered at the lesion (A and B). Data show the means ± SEM (**, p < 0.01; n = 10).

To determine if GCs directly affect ependymal glia, we investigated Nr3c1 expression after exposure to Dex (Fig. 7 A-C). Similar to non-injured controls (Fig. 1), Nr3c1 localized to the nucleus of Gfap-positive ependymal glia in Dex-treated non-injured fish. However, at 24 and 48 h after spinal cord transection, Nr3c1 persisted in Dex-treated fish, and was localized in the nucleus of ependymal glia consistent with elevated receptor activity. The maintenance of nuclear Nr3c1 immunoreactivity in ependymal glia demonstrates that GCs target these cells.

**Figure 7:**
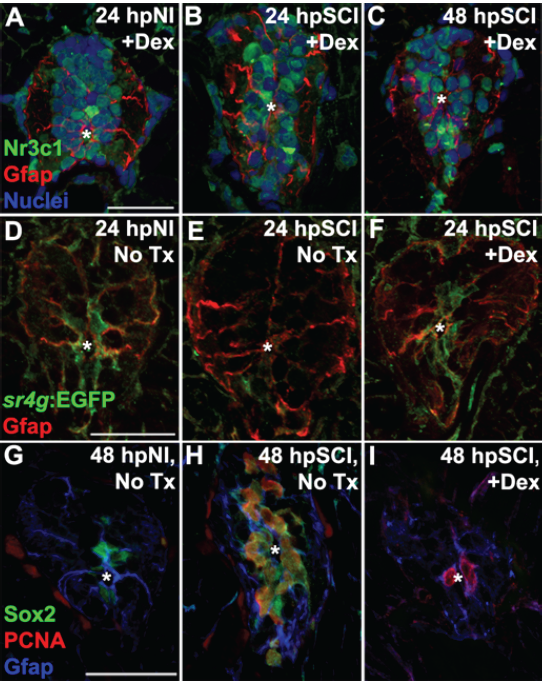
Direct Effects of Glucocorticoids in Ependymal Glia. (A-I) Zebrafish spinal cord transverse sections. (A-C) Immunolabeling of wild-types for Nr3c1 (green), Gfap (red), and DAPI staining (blue) with Dex treatment at 24 h post no injury (A), 24 h (B) and 48 h post SCI (C). Nuclear Nr3c1 (active) localization in Gfap-positive cells around the central canal is stimulated with Dex treatment. See supporting Fig. S1. (D-F) Immunolabeling for EGFP in SR4G transgenic reporter line (green) and Gfap (red) at 24 h post NI (D) and SCI with no treatment (E) or 24 h post SCI +Dex (F). Nr3c1 signaling is constitutively active without injury (D), diminished after SCI (E), and maintained after SCI + Dex. (G-I) Immunolabeling for Sox2 (green), PCNA (red) and Gfap (blue) from 48 h post no injury (G) or 48 h post SCI followed by no treatment (H) or 48 h post SCI +Dex (I). Expression of Sox2 in ependymal glia and NPCs is downregulated after Dex treatment. Representative images from n = 10 spinal cords. Scale bars, 20 µm. A is the same as B, C; D is the same as E, F; G is the same as H, I. * denotes central canal.

To further identify the GC-responsive cell-types relevant to repairing the transected spinal cord, we employed the SR4G transgenic zebrafish line to report GC receptor signaling (46); Gfap immunostaining identified ependymal glia. GC receptor signaling colocalized in non-injured control tissue with ependymal glia surrounding the central canal. In agreement with our Nr3c1 analyses, the intensity of EGFP expression in ependymal glia in juxta-lesional cross sections of cord tissues was reduced 24 h after transection, but was retained in the transected cords of Dex-treated fish (Fig. 7 D-F). These observations support our hypothesis that GCs are an inhibitory cue for regeneration after cord transection. But in contrast to previous studies (21, 22), our findings have established a new paradigm, that GCs elicit inhibitory effects directly on ependymal glia.

A plausible mechanism for GC regulation of ependymal glia is via suppression of injury-induced factors that promote neural repair. Expression of the transcription factor Sex-determining region Y-box 2 (Sox2), a critical component of neural regeneration, increases in zebrafish ependymal glia following spinal cord injury and is required for maximal ependymal glial proliferation (47–49). To determine if GCs regulate Sox2 expression, we administered Dex to maintain high levels of GC receptor signaling in ependymal glia (Fig. 7 D-F). In agreement with previous studies, ependymal glia of non-injured controls and daughter neural precursor cells at the peak of proliferation (48 h after transection) expressed Sox2 (Fig. 7 G, H). Dex treatment resulted in reduced Sox2 expression in both quiescent ependymal glia and dividing neural precursor cells at 48 h after transection. These results indicate that GC receptor signaling in spinal cord regeneration occurs upstream of Sox2, and suggest that GCs act on ependymal glia to suppress its expression and inhibit neural repair.

## Discussion

Our study compares responses to spinal cord injury in larval zebrafish and rats. An important discovery is that the glucocorticoid receptor Nr3c1 in ependymal glia is a negative regulator of functional regeneration and is differentially regulated in the two species. By directly targeting ependymal glia, GCs block functional recovery by inhibiting cell-cycle re-entry and formation of trans-lesional glial bridges. Thus the mammalian spinal cord’s limited capacity to regenerate could be plausibly explained by maladaptive NR3C1 regulation in ependymal glia during acute phase spinal cord injury. Our data question the rationale for use of high-dose corticosteroid therapy in the clinical management of acute traumatic spinal cord injuries.

Ependymal glia are of central importance in the zebrafish spinal cord’s ability to regenerate. In response to injury, these cells re-enter the cell cycle and give rise to multipotent neural progenitor daughter cells that replace lost neural cells and form bipolar trans-lesional bridges to support axon regrowth (15, 17–20, 35, 37). Infiltration of inflammatory cells is an early cellular response to CNS injury in both mammals and zebrafish (50–52). The literature notes that administration of the glucocorticoid Dex as an immunosuppressive agent following traumatic injury in the adult zebrafish brain (21) or larval spinal cord (22) counteracts hematogenous cells and microglia in injured tissues, upregulation of pro-inflammatory cytokines, and neural regeneration. Based on these reports, we hypothesized that GCs were inhibitory to regeneration. We sought to clarify the mechanism by identifying and comparing the cellular targets of GCs administered after cord transection in larval zebrafish and rats.

Our finding of Nr3c1 immunolocalization throughout spinal cord tissues in both larval zebrafish and adult rats supports wide-spread expression in the CNS. Whereas the nuclear Nr3c1 localization in ependymal glia of non-injured fish indicates intrinsic signaling activity, the cytoplasmic distribution in rats suggests low constitutive activity without injury. The changes in Nr3c1 distribution induced by spinal cord injury in the two species supports the notion that signaling activity is regulated bi-directionally: becoming cytoplasmic (inactive) in zebrafish and nuclear (active) in rats. Investigation in the SR4G transgenic reporter zebrafish confirmed that GC receptor signaling was constitutively active in ependymal glia, but it was less active 4 h after injury (not shown) and was no longer detectable at 24 h, coinciding with the onset of cell proliferation and glial bridge formation. Most convincing in this regard was the observation that in fish treated with Dex after cord transection, Nr3c1 persisted in the nucleus of ependymal glia and GC receptor signaling continued. These data established ependymal glia as novel targets of GCs. The accompanying reduction of Sox2 expression in both ependymal glia and neural progenitors further supports our hypothesis and identifies GCs as an early determinant of functional regeneration.

GC receptor stimulation by the agonist Dex did not alter locomotor responses in control zebrafish but was sufficient to block recovery after cord injury. Reversal of this blockade by TALEN-mediated mutagenesis of *nr3c1* indicates that locomotor recovery is Nr3c1-dependent. Anticipating that interdependent cellular events are initiated by spinal cord injury, we investigated the effect of GCs at defined time points in the course of regeneration. GC administration did not alter cell viability, but the resulting inhibition of ependymal glial proliferation may have contributed to the limited extent of neurogenesis. Similarly, the GC-induced blockade of trans-lesional glial bridge formation in turn may have reduced axon regeneration. Taken together, these data reveal that GCs are inhibitory to pro-regenerative properties in ependymal glia. However, we cannot rule out direct effects of GCs on additional regenerative processes by non-mutually exclusive mechanisms.

The mechanisms underlying GC regulation of ependymal glia are poorly understood. Secretion of pro-inflammatory factors by recruitment of hematogenous cells and microglia has been implicated as a stimulus for regeneration. Our finding that regeneration was initiated in the ‘bloodless’ *cloche* mutant zebrafish (39, 45) argues against this hypothesis, and suggests that neural signaling is sufficient to stimulate ependymal glia to reenter the cell cycle.

The data presented here justify a revised model of GC signaling following CNS injury that includes activation of an inhibitory signaling cascade (via Nr3c1) within radial glia. The present study demonstrates that ependymal glia in zebrafish have constitutive Nr3c1 signaling activity, which is down-regulated following cord injury. In addition to their immunosuppressive effects (21, 22), GCs limit the pro-regenerative response of ependymal glia via Nr3c1 stimulation. Regenerative activity occurring in the immune-deficient *cloche* background favors a direct action of GCs on ependymal glia as the basis for regenerative failure.

A direct effect of GCs on ependymal glia was unanticipated in light of the prevailing concept that GCs act by suppressing hematogenous and microglial stimuli to trigger radial glial proliferation in injured adult zebrafish brains (21) and larval spinal cords (22). However, GC receptor expression and signaling activity were not examined in previous studies, leading to the assumption that Dex acted solely as an immune suppressant. Furthermore, immune-deficient zebrafish lines were not used. It remains to be determined whether or not regulatory mechanisms operating in the adult brain (21) differ from those in larval spinal cord tissues. Mechanical lesioning of the brain or sterile inflammation triggered radial glia proliferation and neurogenesis and correlated with inflammatory cell infiltration and expression of pro-inflammatory cytokine mRNAs, including *tumor necrosis factor* α (*tnfα*). These outcomes were attenuated by Dex. The cellular sources of pro-inflammatory/putative pro-regenerative signals were not determined. Mechanistic studies of retinal regeneration in zebrafish reveal that Tnfα is highly expressed upon injury, not by immune cells, but by dying retinal neurons and, subsequently, radial Müller glia. Tnfα release from dying neurons is required to activate regeneration-promoting transcription factors in Müller glia and stimulate their proliferation (53). In the absence of injury, Tnfα in combination with inhibition of Notch signaling is sufficient to stimulate Müller glial proliferation, production of multipotent neural progenitor cells and new neurons (54). Interestingly, Kyritsis et al. (21) detected Cysteinyl leukotriene receptor 1 (CysLT1) in radial glia and showed that its pharmacological blockade attenuated neural repair in the brain. These findings concur with the retinal regeneration findings (53, 54) and implicate pro-inflammatory signaling factors as an initiator of CNS regeneration. Our results from an immune-deficient *cloche* background suggests that the signals are not necessarily of hematogenous or microglial origin. This revised model recognizes inhibitory Nr3c1 signaling by ependymal glia and accounts for direct effects of GCs after injury.

GC therapies are commonly used in treating acute spinal cord injuries to limit damage by dampening inflammation (1, 2). Despite lack of supportive evidence for this clinical practice and systemic complications of GC therapy (5–8), the practice continues (3). The purpose of this study was to determine the role and mechanism of GCs during regeneration after transecting spinal cord injury. We have identified NR3C1 as plausibly a maladaptively regulated factor that in turn may contribute to the irreversible paralysis in mammals. This new evidence advocates a reexamination of experimental and clinical strategies to determine if anti-GC therapies, with or without synergistic agents, might improve functional outcomes. For example, administration of recombinant FGF following spinal cord injury in rodents supported modest functional improvement (55, 56) and stimulated formation of peri-lesional polarized astroglial structures (57). Unknown inhibitory mechanisms in mammals likely block trans-lesional glial bridges and robust functional recovery regardless of putative stimulatory factors. The functional benefit of GC receptor suppression merits investigation.

## Acknowledgements

This work was supported by the National Institutes of Health (NS67311, JRH); a John M. Nasseff, Sr. Career Development Award in Neurologic Surgery Research from the Mayo Clinic (JRH); the National Institute of Biomedical Imaging and Bioengineering (EB02390 and 2T32EB005583, AJW); National Center for Advancing Translational Sciences (TL1 TR002380, AJW); Mayo Foundation, Bowen Foundation, Kipnis fund for Regenerative Medicine (AJW). We appreciate the assistance of the Mayo Clinic zebrafish core facility technicians, rodent core facility technicians for care provided to animals, the Mayo Microscopy and Cell Analysis Core for experimental and technical support. We thank Jarred Nesbitt for assistance in zebrafish husbandry and post-operative care of rats, Stephen C. Ekker, Ph.D. for thoughtful discussions while preparing the manuscript, David R. Hyde Ph.D. for providing transgenic zebrafish lines and Bingkun Chen Ph.D. for performing rat surgeries and training.

## Author Contributions

Conceptualization, C.M.N. and J.R.H. Methodology, C.M.N, N.N.M., and J.R.H. Validation, C.M.N., H.L., R.G.K., and A.K. Formal Analysis, C.M.N. Investigation, C.M.N., H.L., R.G.K., and A.K. Writing – Original Draft, C.M.N. and J.R.H. Writing – Editing and Reviewing, C.M.N., H.L., R.G.K., N.N.M., K.J.C., V.A.L., A.J.K., and J.R.H. Visualization, C.M.N. Supervision, K.J.C., V.A.L., A.J.K., and J.R.H. Project Administration, J.R.H. Funding Acquisition, J.R.H.

## Supplemental Information

**Supplementary Figure S1:**
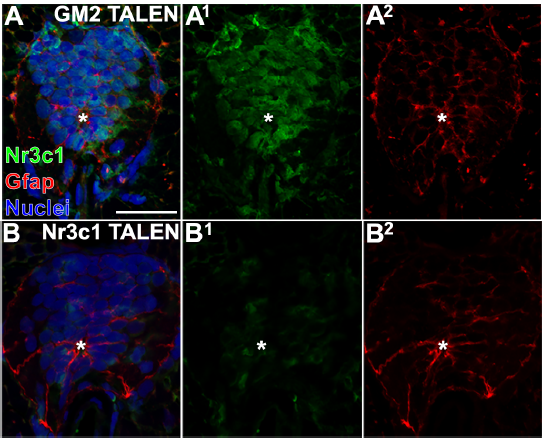
**TALEN-mediated Mutation of *nr3c1***. (A, B) Zebrafish spinal cord transverse sections. Embryos were injected with non-targeting GM2 (A-A^2^) or *nr3c1* targeting TALENs (B-B^2^), then raised for 4 dpf and show immunolabeling for Nr3c1 (green), Gfap (red) and DAPI staining. Scale bar, 20 µm (A is the same as B). * denotes the central canal. Representative images from *n* = 10 spinal cords. The data support Fig. 2.

**Supplementary Figure S2:**
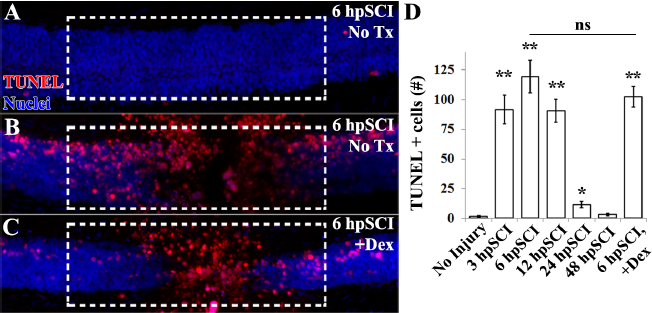
Glucocorticoids do not Alter Cell Viability Following SCI. (A-C) Whole-mount preparations (lateral view) of wild-type larval zebrafish show detection of TUNEL (red) and DAPI staining (blue) in the spinal cords from no injury and no treatment (A), and 6 h post SCI with either no treatment (B) or +Dex (C). Dashed boxes represent a 200-µm region centered at the lesion site used for quantifications. Quantification of the mean number of TUNEL-positive nuclei at 3-, 6-, 12-, 24-h post SCI and 6 h post SCI +Dex (± SEM). \**p* < 0.05, ** *p* < 0.01 (compared to no injury); ns = not significant. *n* = 10.

**Supplementary Figure S3:**
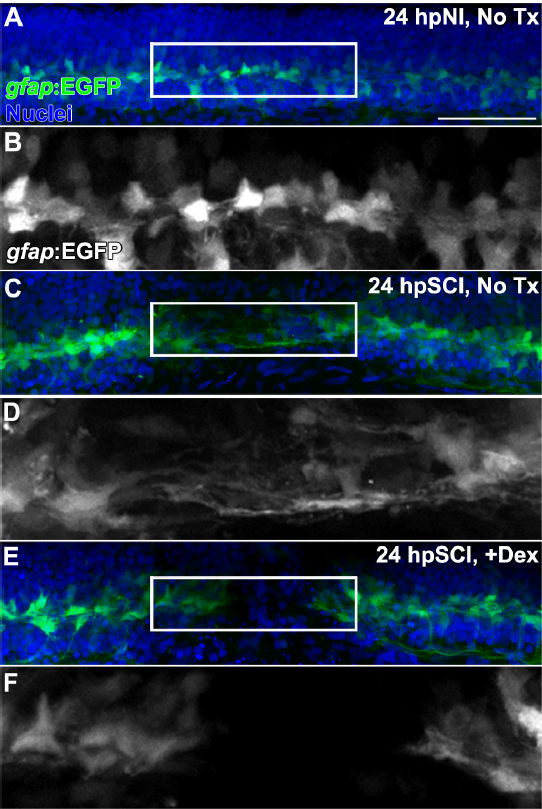
Glucocorticoids Inhibit Formation of Trans-lesion Glial Bridges. (A-F) Whole-mount preperations (lateral view) of Tg(*gfap*:EGFP) transgenic zebrafish show immunlableing for EGFP (green) and DAPI staining (blue) from 24 h post no injury (A and B), 24 h post SCI with no treatment (C and D) and 24 h post SCI +Dex (E and F). EGFP-positive ependymal glia were observed spanning the lesion site at 24 h post SCI in controls (C and D), and were absent with Dex (E and F). Scale bar, 50 µm. Boxed regions (A, C, E) are magnified by 175% (B, D, F, respectively). Representative images from n = 10. See complementary Fig. 4.

## STAR Methods

### Contact for Reagent and Resource Sharing

Further information and requests for resources and reagents should be directed to and will be fulfilled by John R. Henley, Ph.D.

### Experimental Model and Subject Details

Wild-type stocks were derived from offspring of adult zebrafish purchased from Segrest Farms (Segrest Farms, Florida, USA). Studies were approved by the Mayo Clinic Institutional Animal Care and Use Committee (IACUC protocol A36315-15). Care of fish accorded with standard protocols. Adult and embryonic fish were maintained at 28.5 °C on a 14-h light/10-h dark cycle. Larvae were raised in standard E2 embryo media on a diet of paramecium starting at day 5 post-fertilization (dpf); brine shrimp supplementation began at 14-28 dpf, and was the sole diet thereafter.

Experiments involving rats were approved by the Mayo Clinic (IACUC protocol A00001766-16). Rats were housed per National Institutes of Health and U.S. Department of Agriculture guidelines on a 12 h light-dark cycle and standard food and water ad libitum. Adult female Fischer rats (total of six, weighing 180-200 g) were used because of ease of handling and less frequent urinary tract infections from bladder squeezing.

## Method Details

### Zebrafish: Spinal cord injury, EdU and Dexamethasone Treatments

Tricaine methanesulfonate (Argent Chemical Laboratories, Redmond, WA) was added to medium in 100 ×; 15 mm Petri dishes (Corning, Durham, NC) as anaesthetic for spinal manipulations, and anesthetic overdose was used for killing. The cord was transected following established protocols (33, 34), with 3 dpf larval fish mounted on Sylgard silicone elastomer (Dow Corning, Midland, MI) stages cast from microinjection molds (Adaptive Science Tools, Worcester, MA). Borosilicate capillary tubing (World Precision Instruments, Sarasota, FL) was pulled, broken and inserted through the cord under a Zeiss Discovery V12 stereoscope. Rare fish that bent along the body axis due to notochord damage were excluded from study. For GC exposure, the synthetic GCR agonist Dex (Sigma) was added to the holding bath (0.01% total volume methanol vehicle) immediately after injury. Vehicle was added to control media. Solutions were replaced every 48 h. Based on published data (58, 59, 60), we performed a dose range study of 1-100 µM to establish 10 µM as ideal concentration (minimal sufficient to inhibit functional recovery following cord transection). This dose did not affect behavioral responses, gross morphology, or survival in non-injured fish exposed for 120 h.

### Rats: Spinal Cord Injury and Post-operative Care

Rats were monitored and cared for by veterinarians experienced in handling rodents with spinal cord injury. Ibuprofen was added to drinking water 40 mg/L beginning 48 h prior to surgery and thereafter. Rats were randomly assigned to experimental groups for laminectomy alone or laminectomy followed by complete spinal cord transection (T9 vertebral level), according to protocols described in Chen et al., (61). Anesthesia was via intraperitoneal ketamine (80 mg/kg; Fort Dodge Animal Health, Fort Dodge, IA), xylazine (5 mg/kg; Lloyd Laboratories, Shenandoah, IA) and inhaled isoflurane (1.5-2%; Cardinal Health, Dublin, OH) as needed to maintain for surgery. Rats were kept on 37 °C heating pad during surgery and recovery from anesthesia. The spinal cord was cut with a sharp blade (No 11, BD Medical, Batavia, IL), then probed with a microhook to confirm complete transection. Rats were housed individually in low walled cages, and soft chow was provided on the cage floor. Following surgery, intraperitoneal saline was injected (5 mL) and bladders were squeezed twice daily. Urine was examined for signs of infection. Analgesics and antibiotics were given as needed.

### Rat Tissue Preparation

After killing by intraperitoneal injection of 0.4 mL pentobarbital sodium (40 mg/kg; Fort Dodge Animal Health, Fort Dodge, IA), and perfusion with saline wash followed by 4% PFA (Chen et al., 2011), the lesioned spinal cord region and adjacent tissues were removed, and fixed for 72 h in PFA at 4 °C. The spinal cord tissue, and approximately 10 mm of adjacent tissue on each side, was isolated, fixed overnight in 4% PFA at 4 °C, then processed for cryosectioning.

## TUNEL

Terminal deoxynucleotidyl transferase-mediated biotinylated UTP nick end labeling (TUNEL) performed on 4% PFA fixed wholemount zebrafish using the ApoAlert DNA Fragmentation Assay kit (Clontech, Mountain View, CA). Removal of pigment by incubation with 1% H2O2 and 5% formamide in 1x PBS under intense light exposure was followed by tissue permeabilzation with Proteinase K. Enzymatic labeling of fragmented DNA was done according to the manufacturer’s protocol in label buffer containing 2% biotinylated dUTP (Roche Diagnostics, Indianapolis, IN). Samples were processed and dUTP was detected by Alexa-fluor conjugated Strepavidin (Life Technologies, Eugene, OR).

### Zebrafish Whole mount Immunohistochemistry and EdU

Zebrafish were fixed overnight at 4°C in either 9:1 ethanolic formaldehyde (100% ethanol:37% formaldehyde; for PCNA) or 4% PFA (4% paraformaldehyde, 5% sucrose, 1x PBS) then stored in methanol at −20°C. Rehydration with 1x PBS containing 1% Triton X-100 and by bleaching with 1% H2O2 and 5% formamide diluted in 1x PBS under intense light exposure was followed by proteinase K permeabilization. For EdU uptake, 0.5 mM EdU was added to E2 immediately following transecting injury. Detection of EdU was performed with 2 h incubation with Click-it EdU AF 594 kit following manufacturer’s protocol (Thermofischer). Incubation in blocking buffer containing 10% normal goat serum, 2% DMSO and 1% Triton X-100 was for 2 hrs at 25 °C. Primary antibodies were diluted in blocking buffer for overnight (up to 48 h) incubation at 4°C with gentle rocking. Primary antibodies used in this study were mouse anti-PCNA monoclonal antibody (1:500; clone PC10, Sigma-Aldrich, St. Louis, MO), rabbit anti-EGFP polyclonal antiserum (1:250; Life technologies, Eugene, OR), chicken anti-GFP polyclonal antiserum (1:500; Abcam, Cambridge, MA) mouse anti-Acetylated tubulin monoclonal antibody (1:200; Sigma Aldrich), mouse anti-HuC/D monoclonal antibody (1:300; Life Technologies). Samples were washed in 1x PBS containing 1% Triton x-100 and incubated in secondary antibody diluted 1:250 in wash buffer overnight at 4°C. Nuclei were labeled with DAPI (1: 5000; ThermoFisher Scientific). The secondary antibodies used in this study were Alexa Fluor goat anti primary IgG 488, 568, 594, and 647 (Life Technologies) then washed in 1x PBS containing 1% Triton X-100 and mounted with glass coverslips and ProLong Diamond antifade reagent (Life technologies).

### Cryopreservation and Cryosection Immunohistochemistry

Zebrafish were fixed in 4% PFA or 9:1 ethanol:37% formaldehyde overnight at 4 °C and washed in 5% sucrose/1x PBS and 30% sucrose/1x PBS and incubated in 2:1 O.C.T.: 30% sucrose/1x PBS for 4 h and frozen (−20 °C) in O.C.T. (Sakura Finetek, Torrance, CA) followed by cryosectioning at a thickness of 16 microns. Frozen sections were dehydrated at 55°C for 2 h and stored at −80°C. Slides were thawed for 20 minutes at 55°C and rehydrated in 1X PBS. Antigen retrieval was done with 10 mM sodium citrate (pH 6.0) at 95 °C with 0.1% Tween-20 for 15 minutes. Sections were incubated in 1X PBS/4% normal goat or chicken serum/0.4% Triton X-100/1% DMSO blocking solution for 1 h at room temperature and overnight incubation at 4°C with the primary antibody diluted in blocking buffer. Primary antibodies used in this study were mouse anti-PCNA monoclonal antibody (1:1000, clone PC10, Sigma-Aldrich), rabbit anti-PCNA polyclonal antiserum (1:500, Abcam), rabbit anti-EGFP polyclonal antiserum (1:750, ThermoFisher Scientific, Waltham, MA), chicken anti-GFP polyclonal antiserum (1:250; Abcam), rabbit anti-GFAP polyclonal antiserum (1:500, Dako, Carpinteria, CA), mouse anti-GFAP (1:250, Sigma-Aldrich), goat anti-Sox2 polyclonal antiserum (1:100, R&D Systems, Minneapolis, MN), Chicken anti-Vimentin (1:400. EMD Millipore), Rabbit anti-Nr3c1 (1:50, ThermoFischer Scientific). Washes were done in 0.05% Tween-20/1x PBS and incubation in secondary antibodies diluted in 0.05% Tween-20/1x PBS for 1 h at 25 °C. Secondary antibodies used in this study were Alexa-Fluor goat or chicken anti-primary IgG 488, 568, and 647 (1:500, Life Technologies, Eugene OR). Nuclei were labeled with DAPI (1:15000; ThermoFisher Scientific). Washes were repeated and mounted with glass coverslips and Prolong Diamond Antifade Mountant (Life Technologies).

### Behavioral Assays

Locomotor function was scored on a scale from 1 to 5 to measure functional recovery at designated time points. Locomotor recovery assay scores (17) were adapted for larval fish. Scores were; 1, body caudal to the lesion was paralyzed and the fish lay on its side at the bottom of the dish and was non-responsive to tail prod; 2, fish were oriented upright and had no response or responded to tail prod with brief non-productive body movement rostral to the lesion site while areas caudal to the lesion site were paralyzed; 3, tail prod evoked brief uncoordinated movements which produced a short period of locomotion; 4, fish were able to escape and movements were sustained for a longer period of time and became coordinated; 5, locomotion of the spinal cord injured zebrafish were indistinguishable from no injury counterparts. Scores were obtained from the average values of each fish in triplicate.

## TALENs

Targeted mutation of *nr3c1* with Transcription activator-like effector nucleases (TALENs) was performed as previously described in Krug et al (46). Briefly, one-cell embryos were microinjected with 50 pg of mRNA encoding *nr3c1* targeting or non-targeting GM2 TALEN pairs. Mutation of nr3c1 was confirmed by RT-PCR (not shown) and immunohistochemical detection of Nr3c1 (Supp. Fig. 1).

### Quantification and Statistical Analysis

Microscopy was performed with an Olympus Fluoview FV1000 confocal microscope. Cell counts from lateral views of whole-mounted zebrafish were obtained within 100 µm of the center of the lesioned tissue from 20 µm thick confocal z-stacks. Alternatively, 10 µm thick z-stacks were obtained from 6-consecutive spinal cord cross sections that spanned the lesioned tissue (Fig. 3). Between 6 and 10 spinal cords were examined in each experimental time point, actual numbers are reported in figure legends. Images were cropped, high backgrounds reduced and low-intensity signals enhanced in Adobe Photoshop (Adobe Systems, San Jose, CA). Fluorescence levels were modified identically in all layers within a panel and to all other panels in a figure. Statistical significance was calculated by using a Student’s t-test (two-tailed and two-sample unequal variance). Actual p-values are listed in the text, less than 0.05 and 0.01 were considered to be significant and highly significant, respectively. Regeneration of axons was quantified by corrected total fluorescence measurements (38). Integrated density calculations of immunofluorescence were obtained from n= 7 spinal cord tissues that were within 50 µm of the center of lesion using ImageJ software. Measurements for each image were normalized to regions within the same field of view containing background signal. Corrected total fluorescence values were obtained and plotted using Excel software by subtracting the mean integrated density of background measurements from the fluorescence measured in the injury region of spinal cord tissues.

## KEY RESOURCES TABLE

| REAGENT or RESOURCE | SOURCE | IDENTIFIER |
| --- | --- | --- |
| **Antibodies** |  |  |
| Rabbit polyclonal Glial Fibrillary Acidic Protein (GFAP) antibody | Dako | Cat# M0761, RRID:AB_210995 |
| Mouse Anti-Proliferating Cell Nuclear Antigen Monoclonal Antibody, Unconjugated, Clone PC 10 | Sigma-Aldrich | Cat# P8825, RRID:AB_477413 |
| Rabbit Anti-Green Fluorescent Protein (GFP) Polyclonal Antibody, Unconjugated | Molecular Probes | Cat# A-6455, RRID:AB_221570 |
| Chicken Anti-GFP antibody | Abcam | Cat# ab13970, RRID:AB_300798 |
| Monoclonal Anti-Acetylated Tubulin antibody produced in mouse | Sigma-Aldrich | Cat# T7451, RRID:AB_609894 |
| Mouse Anti-Human HuC / HuD neuronal protein Monoclonal Antibody, Unconjugated, Clone 16A11 | Molecular Probes | Cat# A-21271, RRID:AB_221448 |
| Rabbit Anti- PCNA antibody | Abcam | Cat# ab19166, RRID:AB_444779 |
| Monoclonal Anti-Glial Fibrillary Acidic Protein (GFAP) antibody produced in mouse | Sigma-Aldrich | Cat# G3893, RRID:AB_477010 |
| Goat Anti-Human Sox2 Polyclonal antibody, Unconjugated | R and D Systems | Cat# AF2018, RRID:AB_355110 |
| Chicken Anti-Vimentin antibody | Millipore | Cat# AB5733, RRID:AB_11212377 |
| Anti-NR3C1 antibody produced in rabbit | Sigma-Aldrich | Cat# SAB1408583, RRID:AB_10742538 |
| **Chemicals, Peptides, and Recombinant Proteins** |  |  |
| Dexamethasone | Sigma-Aldrich | D1756 |
| DAPI (4',6-diamidino-2-phenylindole) | ThermoFischer | D1306 |
| ProLong Diamond Antifade Mountant | ThermoFischer | P36961 |
| Biotinylated dUTP (Bio-16-dUTP) | Sigma-Aldrich | 11093070910 |
| **Critical Commercial Assays** |  |  |
| ApoAlert DNA Fragmentation Assay kit | Clontech | Cat# 630108 |
| Click-it EdU AF 594 kit | Thermofischer | Cat# C10339 |
| Experimental Models: Organisms/Strains |  |  |
| Zebrafish: Segrest Wild-type | Segrest Farms | N/A |
| Zebrafish: Tg(*gfap*:EGFP)nt11 | Kassen et al., 2007 | ZFIN ID: ZDB-FISH-150901-26596 |
| Zebrafish: SR4G transgenic reporter line | Krug et al., 2014 | ZFIN ID: ZDB-FISH-151006-4 |
| Recombinant DNA |  |  |
| *nr3c1* TALEN pairs | Krug et al., 2014 | ZFIN ID: ZDB-TALEN-150928-1 |
| GM2 TALEN pairs | Krug et al., 2014 | N/A |

